# NITRITE INCREASES MITOFUSIN-1 LEVELS TO INHIBIT VASCULAR SMOOTH MUSCLE CELL PROLIFERATION AND PREVENT INTIMAL HYPERPLASIA

**DOI:** 10.64898/2026.01.27.702091

**Authors:** Wenxi An, Christopher Reyes, Krithika Rao, Cristina Espinosa-Diez, Parth Patel, Danielle Guimaraes, Morgan Jessup, Claudette St Croix, Delphine Gomez, Sruti Shiva

## Abstract

Vascular disease remains a leading cause of morbidity and mortality and is driven by maladaptive vascular remodeling following injury. Stent-induced vascular injury induces vascular smooth muscle cell (VSMC) phenotypic switching from a contractile to a proliferative state, resulting in intimal hyperplasia (IH), restenosis and compromised vessel function. Nitrite, an endogenous oxidation product of nitric oxide and a dietary constituent, attenuates IH after vascular injury; however, its underlying mechanisms remain incompletely understood. Nitrite is known to modulate mitochondrial structure and function, and dysregulated mitochondrial dynamics have independently been implicated in VSMC proliferation. We therefore hypothesized that nitrite attenuates IH by modulating mitochondrial dynamics to suppress VSMC proliferation. Using rat aortic smooth muscle cells (RASMCs), we demonstrate that nitrite treatment inhibits cell cycle progression and cell proliferation through upregulation of mitofusin-1 (Mfn1), a GTPase that catalyzes mitochondrial fusion. Mechanistically, nitrite increased Mfn1 protein levels by inhibiting Mfn1 proteasomal degradation. Mfn1 deletion resulted in enhanced proliferation, loss of contractile gene expression, and decreased expression of antioxidant enzymes including catalase and glutathione peroxidase. Restoration of cellular antioxidant capacity significantly attenuated proliferation and preserved contractile gene expression in Mfn1-deficient cells. Smooth muscle cell–specific Mfn1 knockout mice subjected to carotid artery ligation injury exhibited exacerbated IH compared to wildtype mice. Nitrite administration significantly decreased IH in wildtype mice but not Mfn1-deficient mice. These findings identify endogenous Mfn1 as a critical regulator of VSMC cell cycle progression and as an essential mediator of the vasoprotective effects of nitrite.

## INTRODUCTION

Stent-induced vascular injury initiates a cascade of maladaptive remodeling events that ultimately compromise vessel structure and function through restenosis. A key feature of this response is intimal hyperplasia, characterized by increased cell mass and number in the intimal layer of the vessel wall, which leads to the narrowing of the vascular lumen and disrupted blood flow. While intimal hyperplasia is a multifactorial process involving inflammatory signaling, platelet activation, changes in the extracellular matrix, and endothelial cell dysfunction and apoptosis, vascular smooth muscle cells (VSMC) play a central role in its pathogenesis.^1-3^ Under physiological conditions, VSMC maintain a quiescent, contractile phenotype critical for vessel tone. However, following injury, VSMC undergo a phenotypic switch in which they lose expression of contractile genes and transition to a proliferative, migratory state that drives neointimal formation.^4-6^ Detailed lineage tracing studies and single cell RNA analysis provide compelling evidence that these phenotypic transitions occur in vivo and are important in progression of a majority of vascular diseases.^7-9^ Thus, strategies that prevent this transition are of therapeutic interest for limiting pathological remodeling.

At a molecular level VSMC phenotypic change is a complex process regulated by a network of signals that govern the transition from the contractile to synthetic phenotype. This process involves the coordinated downregulation of contractile markers such as Transgelin (TAGLN), α-smooth muscle actin (ACTA2), and smooth muscle myosin heavy chain (MYH11), with simultaneous activation of proliferative, migratory and synthetic pathways.^10^,^11^ At the gene expression level, transcriptional regulators such as Krüppel-like factor 4 (KLF4) and ELK-1 promote phenotypic switching by repressing smooth muscle–specific genes, while loss of Myocardin expression, and consequent disruption in its association with–serum response factor is a permissive step to downregulating the contractile phenotype.^16-18^ Reactive oxygen species (ROS) act as second messengers in this context, modulating redox-sensitive signaling pathways and transcription factors that drive phenotypic switching.^12, 13^ Central to this process are growth factor signaling cascades, including platelet-derived growth factor (PDGF), transforming growth factor-β, and fibroblast growth factor, which both stimulate ROS production and promote cell cycle re-entry.^14, 15^

Recent work highlights mitochondrial dynamics as a regulator of VSMC phenotype.^19-21^ Mitochondria continuously undergo fusion and fission events to build and break cellular mitochondrial networks that dynamically modulate mitochondrial function including metabolism, oxidant generation and ATP production.^22, 23^ Mitochondrial dynamics are closely linked to the cell cycle to ensure that the dividing cell can adapt to changing energetic and biosynthetic needs. For example, mitochondrial fission is required during mitosis and cytokinesis to ensure distribution of sufficient mitochondria to daughter cells. In contrast, mitochondrial fusion increases energetic efficiency during G1 and S phase to support growth and DNA replication.^24, 25^ In VSMC, mitochondrial fusion, catalyzed by the small GTPases mitofusins 1 and 2 (Mfn1 & Mfn2), maintains cells in their quiescent contractile phenotype, while mitochondrial fission, catalyzed by dynamin-related protein-1 (Drp1) is associated with VSMC proliferation and migration.^20, 26-28^ Recent studies that modulate these small GTPase proteins have established a causative role for mitochondrial dynamics in altering VSMC phenotype. Inhibition of Drp1 has been shown to attenuate VSMC migration and proliferation ^28^, while the deletion of Mfn2 potentiates proliferation through disruption of mitochondrial-ER calcium regulation.^26^ Notably, few studies have investigated the role of Mfn1, which has higher GTPase activity and distinct functions (beyond mitochondrial fusion) from Mfn2, in cellular homeostasis.^29, 30^ Further, therapeutic strategies that propagate mitochondrial fusion and are translatable to human vascular injury have not been identified.

Nitrite (NO_2_^-^), traditionally considered an inert product of nitric oxide (NO) metabolism, is now recognized as a bioactive reservoir of NO and signaling molecule with important vascular effects. Nitrite is derived from oxidation of endogenous NO as well as dietary sources and functions in high picomolar levels as a vasodilator in the vasculature. ^31-34^ Nitrite has demonstrated significant therapeutic potential in mitigating intimal hyperplasia by inhibiting VSMC proliferation in pre-clinical models.^35-37^ For example, systemic or local delivery of nitrite attenuated arterial wall hyperplasia following vascular injury in rats and dietary supplementation of nitrite attenuated established intimal hyperplasia.^35^ In a clinical trial of patients who underwent percutaneous coronary intervention, dietary nitrite was associated with a reduction in the ratio of carotid intimal to medial arterial thickness and an improvement in in-stent arterial narrowing.^38^ While these studies demonstrate the therapeutic benefit of nitrite in human vascular disease, the exact mechanism of nitrite action is unclear. Prior pre-clinical studies show that this effect is partially dependent on xanthine oxidase mediated reduction of nitrite to NO and associated with the activation of p21.^35^ However, other studies suggest that nitrite action is independent of its reduction to NO.^39^ Further, while it is established that nitrite modulates mitochondrial function and dynamics in other cell types^39-42^, the role of nitrite in modulating mitochondrial fusion to prevent VSMC proliferation has not been considered.

Herein, we hypothesize that nitrite inhibits VSMC proliferation by targeting mitochondrial dynamics. We demonstrate that nitrite inhibits VSMC proliferation through a mechanism dependent on Mfn1. Further, we elucidate a novel mechanism by which the loss of Mfn1 downregulates cellular antioxidant defenses to propagate VSMC proliferation. Using an in vivo model of vascular injury, we show that the loss of Mfn1 exacerbates intimal hyperplasia and nitrite attenuates intimal hyperplasia in a Mfn1-dependent manner. The implications of these results will be discussed in the context of the physiological and pathological role of mitochondrial dynamics and the therapeutic effects of nitrite in vascular disease.

## METHODS

### Cell Culture, transfection and treatment

Rat aortic smooth muscle cells (RASMC) were cultured in growth media (GM) DMEM/F-12 containing 10% FBS or serum-free media (SS) supplemented with 0.2 mM L-ascorbic acid, 5 μg/ml apo-transferrin and 6.25 ng/ml Na-selenite. Cells were cultured for 48h prior to transfection with 50nM siRNA to sramble (Dharmacon D-001810-01-20), Mfn1 (Dharmacon L-099253-02-0010) or March5 (Dharmacon L-085828-02-0005) duplexed with TransIT-X2 (Mirus MIR 6000) in serum-free Opti-MEM. Sodium nitrite (0 – 25 μM; Sigma-Aldrich 563218) and PDGF-BB (10 ng/mL; Sigma-Aldrich P3201) were prepared in sterile water and sequentially added to RASMC culture media for 24 or 48h based on the experiment.

### Cell cycle and proliferation analysis

DNA was isolated from RASMC (1 million cells) cultured in medium containing 3H-thymidine (5 μCi/mL). Treated and labeled cells were used to precipitate DNA using cold 5% TCA, followed by sequential TCA (5%) and PBS washes. Labeled DNA was dissolved in NaOH to measure 3H-thymidine incorporation (Ultima Gold scintillation fluid, Perkin Elmer). Cell cycle analysis was performed on RASMC fixed and incubated in propidium iodide (50 μg/mL). Singlet cells were gated on a LSRFortessa cell analyzer (BD Biosciences) and frequency of G1, S or G2 population was quantified by FlowJo.

### Immunoprecipitation and Western blot

RASMC (500 – 1000 μg of soluble protein extracts) were incubated with antibody against Mfn1 (Abcam 104274) at 4°C overnight. Protein A/G magnetic beads (Pierce) were incubated with the antigen/antibody mixture, and the conjugates were isolated using a magnetic stand. Targets were eluted by glycine extraction and samples were prepared for Western blot analysis. RASMC lysate and immunoprecipitated samples were subject to Western blot as previously described.^39^ Samples were separated on tris-glycine gels followed by wet transfer onto a nitrocellulose membrane for staining with primary and appropriate secondary antibodies (**Supplemental Table 1**). The membranes were scanned using an Odyssey Infrared Imaging System (LI-COR Biosciences).

### Immunofluorescence imaging of RAMSC and blood vessels

RASMC grown and treated on coverslips were fixed in 2-4% paraformaldehyde, permeabilized with Triton X-100 and stained with primary antibodies overnight in donkey serum at 4°C. Following washes, coverslips were incubated with corresponding secondary antibodies (**Supplemental Table 1**) and nuclear counterstain (NucBlue Live ReadyProbes Reagent Thermo Fisher R37605) for 1h at room temperature. Coverslips were mounted (Thermo Fisher P36930) for imaging. 10-micron tissue sections were deparaffinized and underwent antigen retrieval (Vector Labs H-3300) prior to blocking (PBS containing 0.6% fish skin gelatin and 10% horse serum) and overnight incubation with primary antibodies at 4°C. Sections were then washed and stained with Acta2-FITC to label SMC and DAPI (Thermo Fisher R37605) for nuclear staining and mounted with coverslips containing Prolong Gold mounting solution (Invitrogen P36930). Images were acquired at 20x or 60x (oil) on a Nikon A1 Confocal Microscope System or on a Leica, DMi8 fluorescent microscope. Images were processed and analyzed using ImageJ.

### Real-Time PCR

RNA purified from RASMC by phenol chloroform extraction and aqueous phase purification (Qiagen 74104) was used to prepare cDNA (Bio-Rad 1708891). Real-Time PCR was performed (QuantStudio5, Thermo Fischer) to calculate relative abundance of target genes normalized to β-actin or 18S as the internal control (**Supplemental Table 2**). Quantitative RT PCR was performed on total genomic DNA isolated from samples using unique primer sets to mitochondrial and nuclear encoded genes to establish the represnetation of mitochondrial genome as a measure of cellular mitochondrial content.

### Seahorse Extracellular Flux Analysis Assay

20,000 RASMC were plated 96 well culture plate for measurement on a Seahorse XFe96 Extracellular Flux Bioanalyzer (Agilent) in unbuffered DMEM (10 mM D-glucose, 1 mM sodium pyruvate and 2 mM L-glutamine). Oxygen consumption rates under basal conditions, proton leak (oligomycin; μM), maximal uncoupled oxygen consumption (FCCP; 1.5 μM) and by non-mitochondrial processes (rotenone; 2 μM) were measured sequentially and normalized to cell numbers in each condition as previously described. ^39^

### ROS and ATP measurements

50,000 RASMC were incubated with MitoSOX Red (Invitrogen M36008; 10 μM) or Amplex Red (Invitrogen A22188; 10 μM) and immediately subject to a kinetic measurement of fluorescence intensity at 510/580 nm and 530/590 nm, respectively as previously described.^40^ Similarly, ATP levels in RASMC were measured by a luciferin-luciferase coupled assay using an ATP Determination Kit (Invitrogen A22066) according to the manufacturer’s instructions. Fluorescence intensity was measured kinetically at 485/530nm. All data were calculated as slope of the curve over 30-minute assay period.

### Generation of smooth muscle specific MFN1 knockout mice

All animal experiments were carried out according to approved protocols by the University of Pittsburgh IACUC committee. Female B6.129(Cg)-Mfn1^tm2Dcc^/J mice carrying homozygous loxP-flanked Mfn1 alleles (Mfn1^fl/fl^) (Jackson Laboratories) were bred to male B6.FVB-Tg(Myh11-cre/ERT2)1Soff/J mice (*Myh11*-Cre^ERT2^) which carry a tamoxifen-inducible Cre recombinase under the *Myh11* promoter (Jackson Laboratories. Strain 019079) and B6.129X1-Gt(ROSA)26Sor^tm1(EYFP)Cos^/J mice (Jackson Laboratories, strain 006148) to insert YFP lineage tracer into smooth muscle cells with and without Mfn1. YFP lineage tracing mice are previously described in ^17^ Male mice were exclusively used for experiments due to Y chromosome localization of *Myh11*-Cre^ERT2^ transgene. 6 – 8 weeks old mice were injected intraperitoneally over 10 days with 10 mg/mL tamoxifen (Sigma Aldrich T5648-1G) for cre-lox recombination.

### Carotid Ligation Model and Intimal Hyperplasia

Wildtype and SMC-Mfn1^-/-^ mice were subjected to carotid ligation 2 weeks after tamoxifen injection as previously described. ^43^ Briefly, mice were anesthetized, had blunt dissection of their skin and fascia to expose their vasculature. A 7-0 silk suture was looped around the right common carotid artery and achieved complete ligation. After recovery, mice were placed into cages containing either normal drinking or drinking water supplemented with nitrite (1.5 g/L). After 3 weeks, mice were euthanized and tissues perfused and fixed using a gravity perfusion system. Carotid arteries were isolated and embedded into paraffin blocks. Mason Trichrome staining was performed to quantify intimal hyperplasia. Sections were fixed in Bouin’s solution at room temperature overnight followed by nuclear staining with Weigart’s iron hematoxylin, scarlet-acid fuchsin for muscle and aniline blue for collagen staining. Images were obtained using Trinocular Inverted Fluorescence Phase Contrast Microscope using a 20x objective lens (Zeiss) followed by ImageJ quantification.

### Statistical Analysis

Data are represented as mean ± SEM. Analyses were performed with GraphPad Prism 9. One-way or two-way ANOVA with post-hoc Sidak test was used for continuous data when comparing between groups. Tukey test was used when only comparing main effect. Two-tailed unpaired student t-test for variables that are normally distributed was used when comparing between only two groups.

## RESULTS

### Nitrite upregulates Mfn1 to inhibit vascular smooth muscle cell proliferation

To determine the effect of nitrite on vascular smooth muscle cell proliferation, rat aortic smooth muscle cells (RASMC) were pre-treated with nitrite (0 – 25 µM) and then stimulated to proliferate by treatment with PDGF-BB (10 ng/mL). Similar to prior studies^35^, nitrite inhibited PDGF-induced cell proliferation in a concentration-dependent manner (**Figure 1A**). Analysis of the cell cycle in RASMC demonstrated that nitrite (25 µM) increased the population of cells in G1 and decreased the number in S phase (**Figure 1B**). Additionally, nitrite treated cells showed decreased expression of the cell cycle modulator, cyclin D1, which drives cell cycle entry and G1 phase progression^44^, and a slight decrease in the cell senescence suppressor CDK2^45^, which did not reach statistical significance (**Figure 1C**). Taken together, these data suggest nitrite attenuates RASMC proliferation by inhibiting cell cycle progression.

**Figure 1.**
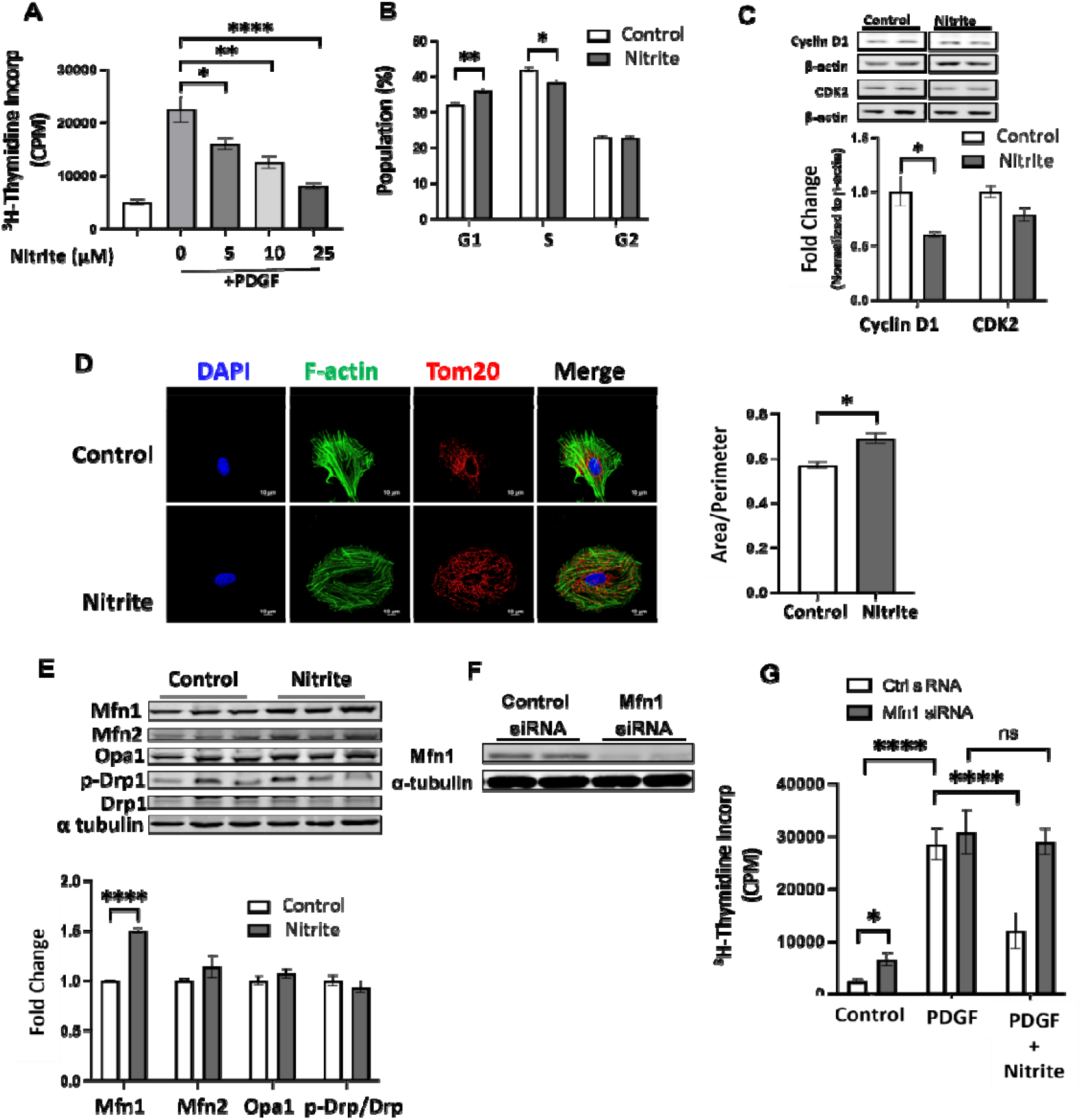
Nitrite inhibits vascular smooth muscle cell proliferation in a Mfn1-dependent manner. **(A)** ^3^H-thymidine incorporation in RASMC pretreated with nitrite (0-25µM) and stimulated with PDGF-BB (10 ng/mL) for 24. **(B)** Percentage of cells in each cell cycle phase in RASMC treated with (black bars) or without (white bars) nitrite (25µM). **(C)** Representative Western blot images (left) and quantification (right) of control (white bars) and nitrite treated (black bars; 25µM; 48 h) RASMC. **(D)** Representative images (left; 60X) of immunofluorescence staining in RASMC treated with or without nitrite (25µM; 24 h). Cells were stained with DAPI to visualize nuclei, for F-actin (cytoskeleton) and Tom20 (mitochondrial membrane). Quantification of fusion (area/perimeter) from 15 cells per replicate is shown on the right. **(E)** Representative western blot (left) and quantification (right) of RASMC treated with (black bars) or without (white bars) nitrite (25µM). **(F)** Representative images of Western blot analysis in RASMC treated with control or Mfn1 targeted siRNA. **(G)** 3H-thymidine incorporation in RASMC RASMC were transfected with control (white bars) or Mfn1 siRNA (black bars) followed by PDGF (10 ng/mL) treatment in the presence or absence of nitrite (25 μM). (n=3); Mean ± SEM; *P<0.05, **P<0.01, ***P<0.001, ****P<0.0001.

Given the known associations between mitochondrial fusion and cell cycle arrest, we assessed the effect of nitrite on mitochondrial dynamics. Visualization of the mitochondrial outer membrane marker Tom20 in RASMCs by confocal microscopy and quantification of mitochondrial area to perimeter ratio revealed elongation of cellular mitochondrial networks after nitrite treatment, indicative of increased fusion (**Figure 1D**). Measurement of proteins that regulate mitochondrial dynamics showed that nitrite significantly increased the protein level of mitofusin-1 (Mfn1), but did not affect levels of Mfn2, Opa1 or phosphorylated Drp1 (**Figure 1E**).

To determine whether the nitrite-mediated inhibition of proliferation was dependent on upregulation of Mfn1, Mfn1 was silenced using siRNA transfection in RASMC (**Figure 1F**). While nitrite inhibited PDGF-dependent RASMC proliferation in control cells, nitrite-mediated inhibition was abolished in cells lacking Mfn1 (**Figure 1G**). Collectively, these data demonstrate that nitrite inhibits RASMC proliferation through a mechanism dependent on Mfn1 upregulation.

### Nitrite increases Mfn1 levels by preventing its ubiquitination

We next examined the mechanism by which nitrite increases Mfn1 expression in RASMC. Nitrite treatment had no significant effect on Mfn1 mRNA levels, indicating that nitrite does not change Mfn1 protein synthesis (**Figure 2A**). To test whether nitrite inhibited Mfn1 degradation, RASMC were treated with or without nitrite in the presence of the protein synthesis inhibitor cycloheximide. While cycloheximide decreased Mfn1 levels in control cells as expected, this effect was significantly attenuated in nitrite treated cells (**Figure 2B**). This data is consistent with a role for nitrite in preventing Mfn1 degradation in RASMC.

**Figure 2.**
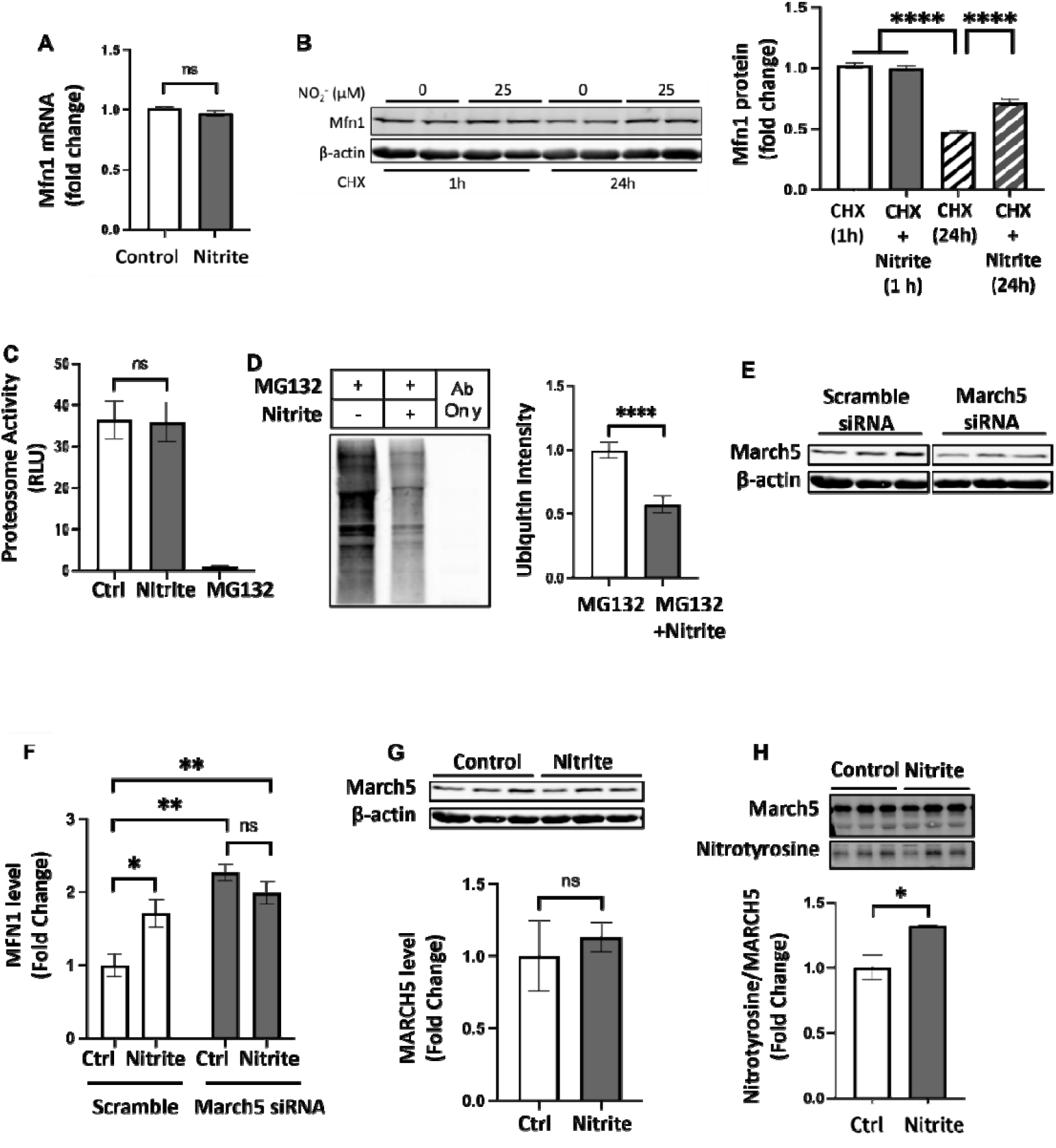
Nitrite increases Mfn1 levels by preventing its ubiquitination. **(A)** Relative Mfn1 mRNA levels in RASMC treated with or without nitrite (n=4). **(B)** Representative images (left) and quantification (right) of Western blots measuring Mfn1 protein levels in RASMC treated with cycloheximide (CHX) in the presence or absence of nitrite (25 μM) for 1 h or 24 h (n=5). **(C)** Proteasomal activity levels in RASMC treated with and without nitrite or the proteosomal inhibitor MG132 (n=5). **(D)** Representative Western blot of ubiquitination of immunoprecipitated Mfn1 (left) and quantification of five independent blots (right). **(E)** Representative images of Western blot analysis in control and March5 knockdown RASMC 48 h after siRNA transfection. **(F)** Representative images (top) and quantification (bottom) of Western blot measuring Mfn1 in RASMC transfected with control or March5 siRNA followed by nitrite treatment (n=5). **(G)** Representative image (top) of Western blot measuring March5 in RASMC treated with or without nitrite and quantification (bottom) of five such blots. **(H)** Representative Western blot of immunoprecipitated Mfn1 from RASMC, stained for March5 and pan-nitrotyrosine (top) and quantification (bottom)(n=5). Mean ± SEM; *P<0.05, **P<0.01, ***P<0.001, ****P<0.**0001**.

Mfn1 is degraded predominantly by the ubiquitin-proteosome system in which Mfn1 is first ubiquitinated by E3 ligases and then degraded by the proteasome^46^. Nitrite treatment did not change the activity of the 20S proteasome in RASMC (**Figure 2C**). However, RASMC treated with MG132 to inhibit proteasomal degradation showed significantly lower levels of Mfn1 ubiquitination with nitrite treatment compared to untreated cells (**Figure 2D**). Given that nitrite inhibited Mfn1 protein ubiquitination, we investigated the role of March5, an E3 ligase which is known to specifically ubiquitinate Mfn1 for proteasomal degradation ^47, 48^ by knocking down March5 with siRNA, which achieved approximately 60% knockdown (**Figure 2E**). In control RASMC, nitrite treatment increased Mfn1 expression (**Figure 2F**). While March5-deficient RASMC showed a basal increase in Mfn1 expression (due to the inhibition of its March5-dependent degradation), there was no further increase in Mfn1 levels after nitrite treatment (**Figure 2F**), suggesting that nitrite increases Mfn1 levels through modulation of March5. Notably, nitrite treatment had no effect on March5 protein levels (**Figure 2G**). However, nitrite did mediate tyrosine nitration of March5, a post-translational modification that potentially inhibits its activity (**Figure 2H**). Taken together, these data demonstrate that nitrite increases Mfn1 protein levels by inhibiting its degradation, potentially through the modulation of March5 activity.

### Deletion of Mfn1 leads to the increased proliferation and decreased expression of contractile genes in RASMC

Since Mfn1 was critical for nitrite-mediated inhibition of proliferation, we next sought to determine the mechanism by which Mfn1 levels regulate RASMC proliferation. Treatment of RASMC with siRNA to target Mfn1 achieved >90% knockdown of Mfn1, with no significant change in the levels of other mitochondrial dynamics proteins Mfn2, Opa1 and Drp1 (**Figure 3A**). While mitochondrial number was not affected by Mfn1 knockdown (**Supplemental Figure 1A**), confocal microscopy examining control and Mfn1 deficient cells demonstrated a decrease in the mitochondrial area/perimeter in

**Figure 3.**
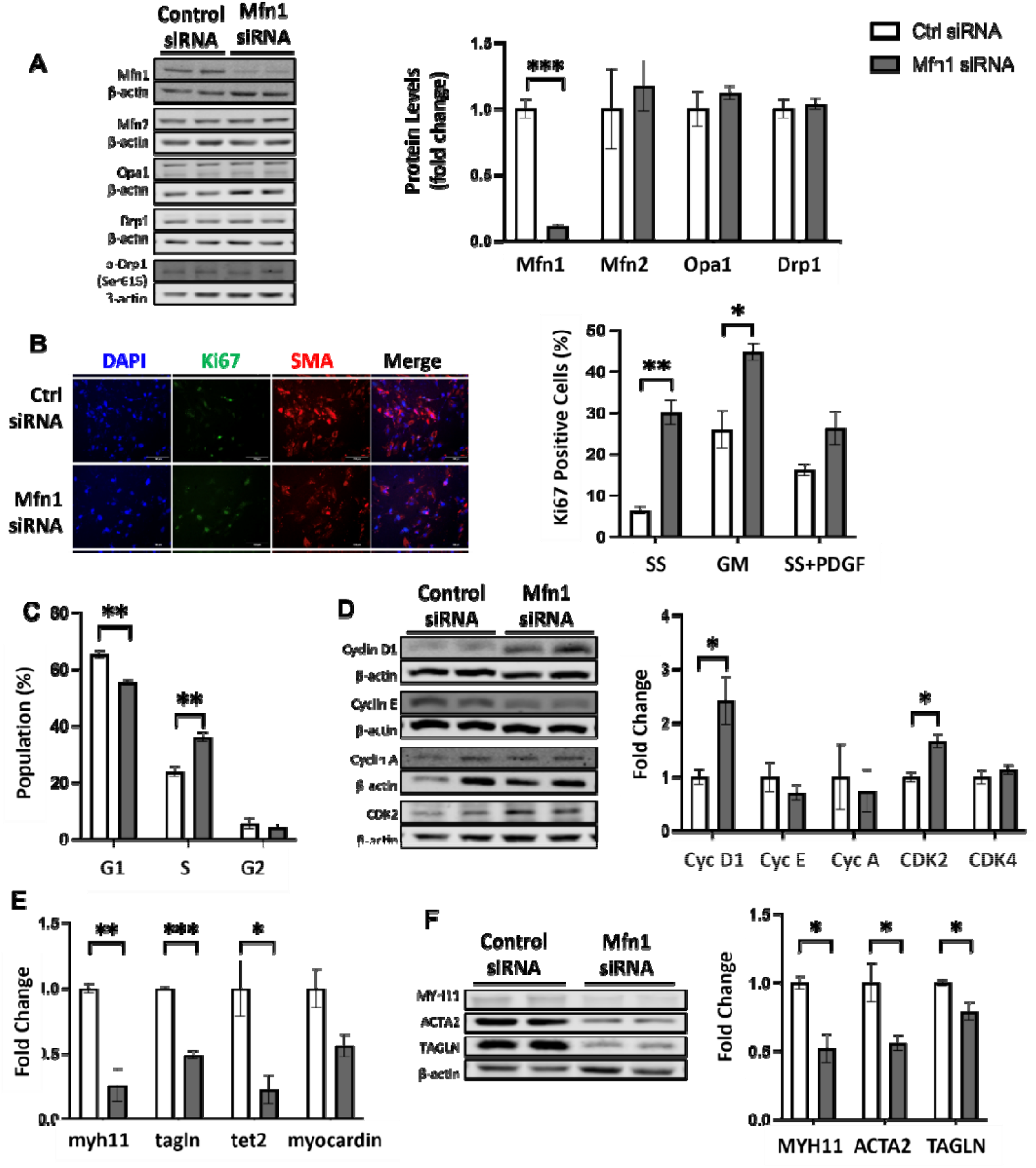
Deletion of Mfn1 leads to the increased proliferation and decreased expression of contractile genes in RASMC. **(A)** Representative Western blot (left) and quantification (right) of protein levels of mitochondrial dynamics proteins (Mfn1, Mfn2, OPA1, phosphorylated Drp1, total Drp1) in RASMC transfected with control (white bars) or Mfn1-targeted (black bars) siRNA. **(B)** Representative images and quantification of immunofluorescence in RASMC transfected with control or Mfn1 siRNA after 48 h in growth media. **(C)** Cell cycle analysis in control (white bars) and Mfn1 knockdown (black bars) RASMC after serum starvation. Quantification shows the percentage of cells present in either G1, S or G2 phase. **(D)** Representative Western blots and quantification of cell cycle regulator protein expression in RASMC transfected with control (white bars) or Mfn1 siRNA (black bars). **(E-F)** Relative change in (E) mRNA levels and (F) protein levels in RASMC treated with control (white bars) or Mfn1-targeted (black bars) siRNA for 48 h. (n=4); Mean ± SEM; *P<0.05, **P<0.01, ***P<0.001, ****P<0.0001.Mean ± SEM; *P<0.05, **P<0.01, ***P<0.001, ****P<0.0001.

Mfn1 deficient cells, consistent with decreased mitochondrial fusion (**Supplemental Figure 1B**). Immunofluorescent staining for Ki67 indicated that Mfn1 deficient cells showed an increase in the number of proliferating cells compared to cells expressing Mfn1. This increase in proliferation occurred with Mfn1 deletion in cells cultured in growth medium, after serum starvation, or with PDGF treatment (**Figure 3B & Supplemental Figure 1C**). Mitofusin-1 deficient cells showed an increased percentage of cells in S phase (**Figure 3C**) and a significant increase in the expression of cyclin D1 and CDK2, consistent with increased cell cycle progression (**Figure 3D**).

To test whether increased RASMC proliferation was associated with a loss of contractility, the expression of contractile phenotypic markers (MYH11, TAGLN, ACTA2, TET2 and myocardin) were measured. Mfn1 deficient RASMC showed a decrease in the mRNA (**Figure 3E**) and protein (**Figure 3F**) expression of contractile phenotypic markers. Collectively, these data demonstrate that Mfn1 deficiency in RASMC leads to proliferation and decreased contractile phenotype.

### Deletion of Mfn1 downregulates cellular ATP production and increases cellular ROS levels

It is well established that mitochondrial fusion potentiates oxidative phosphorylation, which not only increases ATP production but also enhances mitochondrial membrane potential, leading to enhanced physiological mitochondrial ROS (mtROS) production.^22, 40, 49^ Thus, we next characterized the effect of Mfn1 deletion on RASMC mitochondrial function. Extracellular flux analysis showed that Mfn1 deficient RASMC had a significant decrease in basal, maximal and ATP-linked oxygen consumption rate (OCR) (**Figure 4A – B**). Direct measurement of ATP levels showed that Mfn1 deletion decreased ATP levels by 36% ± 9.1% (**Figure 4C**), consistent with decreased oxidative phosphorylation. We next tested whether Mfn1 regulates ROS production. Mfn1 deletion significantly decreased mtROS production by 28% ± 5.5% (**Figure 4D**). Notably, though mtROS was decreased, total cellular hydrogen peroxide (cellular ROS) was significantly increased in Mfn1 knockdown RASMC (**Figure 4D**). To determine the source of the increased cellular ROS in RASMC, the expression levels of non-mitochondrial ROS-producing enzymes (NOX1, NOX4 and lipoxygenase 5 (5-LO)) and antioxidants (catalase, GPX1, Cu/Zn SOD and Mn SOD) were measured. Cells deficient in Mfn1 showed no change in the expression of ROS-producing enzymes, but a significant decrease in the expression of the antioxidants catalase and GPX1 (**Figure 4E**). Collectively, these results demonstrate that the deletion of Mfn1 decreases ATP production and mitochondrial ROS but enhances cellular ROS levels by reducing antioxidant enzyme expression.

**Figure 4.**
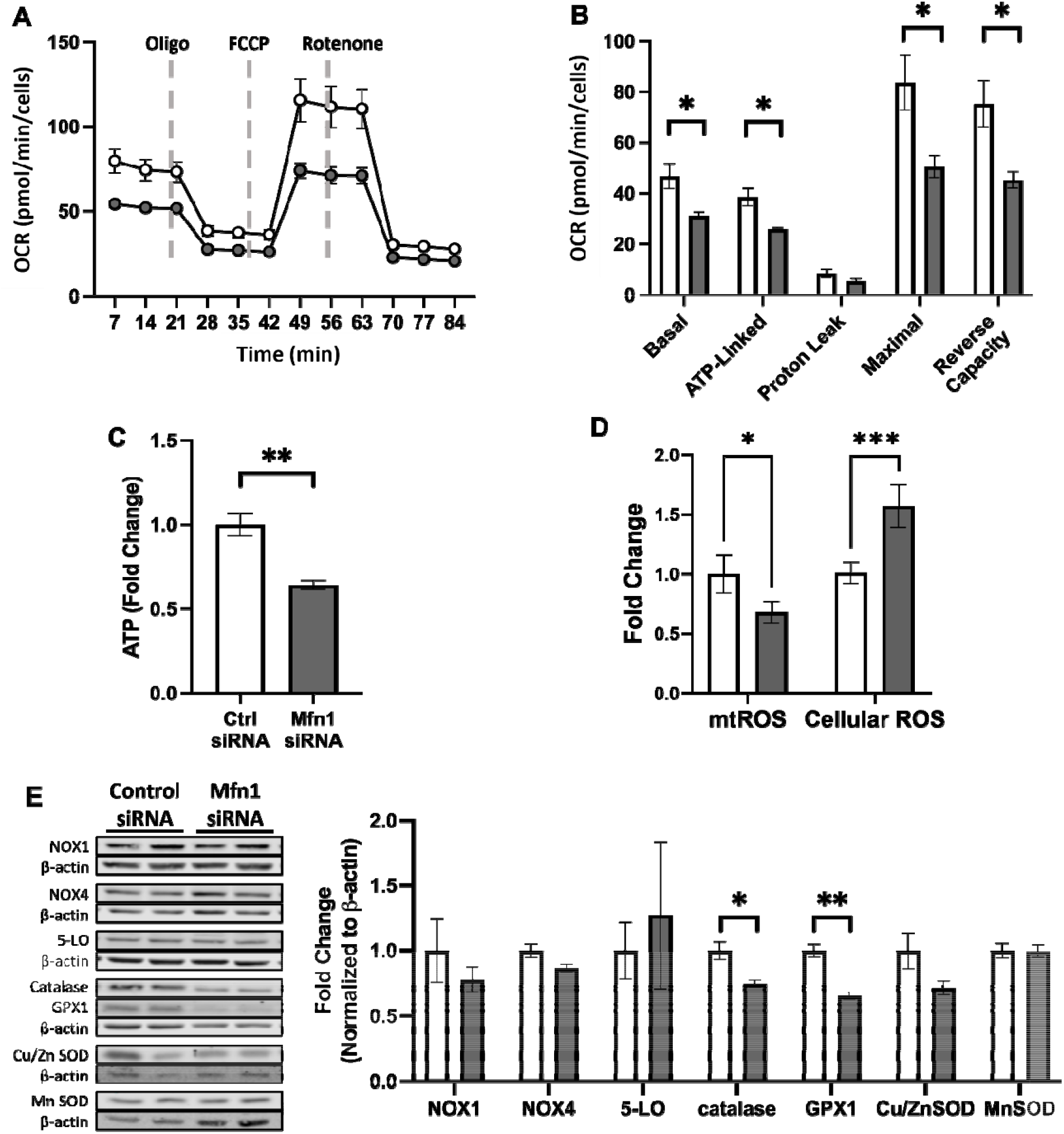
Deletion of Mfn1 downregulates cellular ATP production and increases cellular ROS levels. **(A)** Representative traces and **(B)** quantification of mitochondrial oxygen consumption rate (OCR) and **(C)** relative ATP levels in RASMC transfected with control (white bars) or Mfn1-targeted (black bars) siRNA for 48 h (n=4). **(D)** Fold change in the rate of mitochondrial and total cellular ROS production in RASMC transfected with control (white bars) or Mfn1-targeted (black bars) siRNA for 48 h (n=4). **(E)** Representative Western blot (left) and quantification (right) of protein levels of ROS generating and antioxidant proteins in RASMC transfected with control (white bars) or Mfn1-targeted (black bars) siRNA for 48 h (n=4). Mean ± SEM; *P<0.05, **P<0.01, ***P<0.001, ****P<0.0001.

### Enhanced ROS propagates proliferation in Mfn1 deficient cells

To determine whether increased ROS generation leads to the enhanced proliferation and loss of contractile genes observed with Mfn1 deficiency, adenovirus-induced catalase overexpression was used to scavenge cellular hydrogen peroxide (**Figure 5A**) without affecting ATP levels (**Figure 5B**) in Mfn1 deficient cells. Catalase overexpression restored contractile mRNA (**Figure 5C**) and protein (**Figure 5D**) levels in Mfn1 knockdown RASMC. Catalase overexpression also attenuated the increase in proliferation (**Figure 5E**) in Mfn1 knockdown RASMC. Taken together, these data demonstrate that decreasing ROS generation attenuates the change in proliferation and loss of contractile genes caused by Mfn1 knockdown demonstrating that Mfn1 regulates RASMC function by modulating cellular ROS levels.

**Figure 5.**
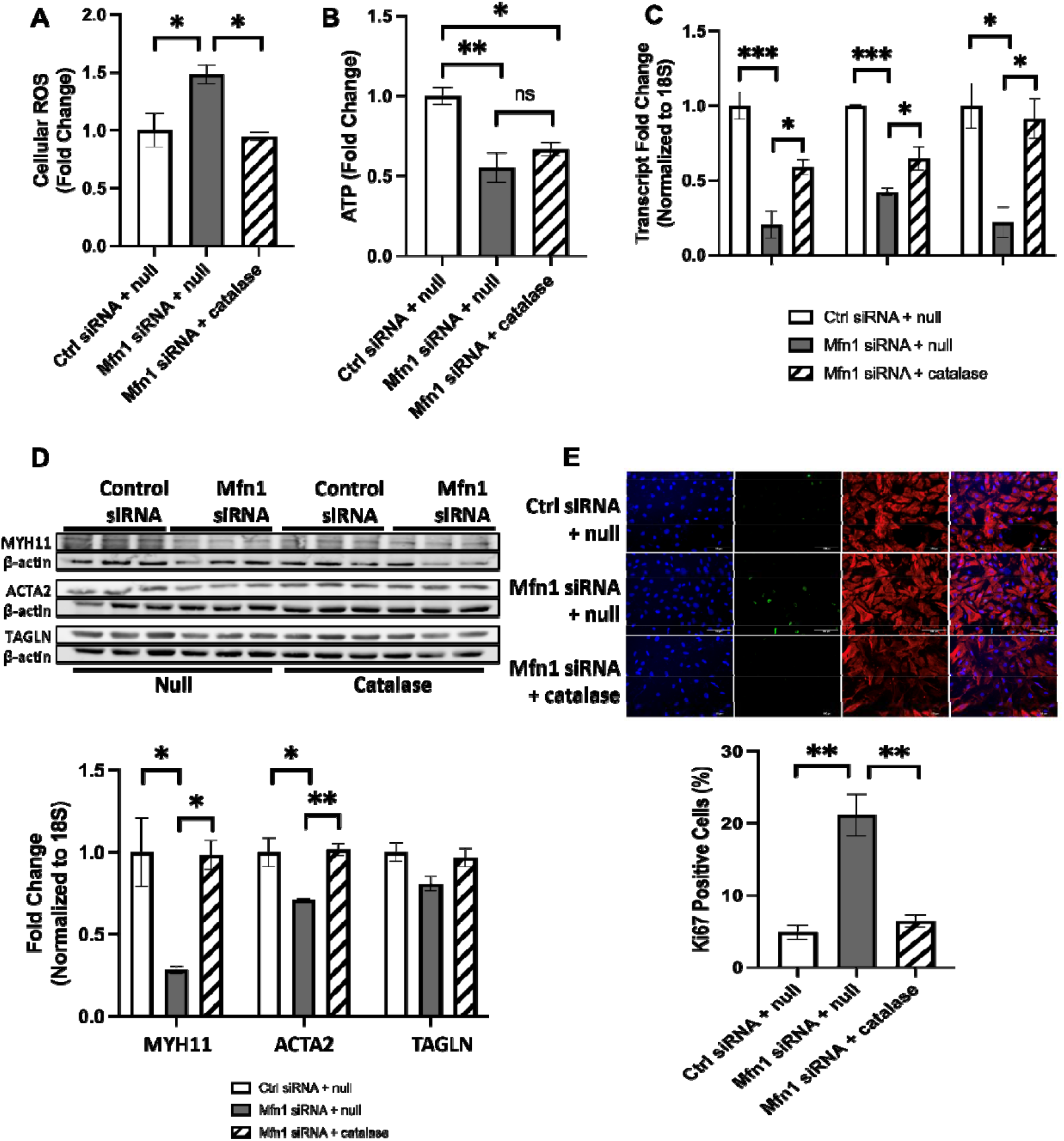
Enhanced ROS propagates proliferation in Mfn1 deficient cells. Relative rates of (**A)** cellular ROS generation and **(B)** ATP production in RASMC transfected with control or Mfn1-targeted siRNA for 24 h followed by empty virus (null) or adenovirus-induced overexpression (2.5×106 PFU/mL) for another 24 h (n=3). **(C)** Relative levels of mRNA for contractile genes in RASMC transfected with control or Mfn1 siRNA for 24 h followed by empty virus (null) or adenovirus-induced overexpression (2.5×106 PFU/mL) (n=3). **(D)** Representative 20X images (top) and quantification (bottom) of immunofluorescence for nuclei (DAPI), proliferation (Ki67), and smooth muscle actin (SMA) (n=3). Mean ± SEM; *P<0.05, **P<0.01, ***P<0.001, ****P<0.0001

### Loss of Mfn1 potentiates vascular remodeling in vivo and nitrite-dependent inhibition of vascular remodeling is dependent on Mfn1 expression

To determine whether Mfn1 prevented smooth muscle proliferation in vivo, SMC-specific Mfn1 knockout mice expressing SMC-specific YFP (SMC-Mfn1^-/-^ mice) were subjected to partial carotid ligation injury to stimulate intimal hyperplasia. Three weeks after vascular injury, histology demonstrated that SMC-Mfn1^-/-^ mice had increased intimal hyperplasia evidenced by higher intima to media ratio, suggesting that SMC-specific Mfn1 knockout exacerbates vascular remodeling (**Figure 6A-B**). Immunofluorescence staining for YFP and Ki67 in the arteries of these mice demonstrated that SMC-Mfn1^-/-^ mice showed significantly more Ki67 and YFP double positive cells, indicative of increased SMC proliferation within the vessel wall after vascular injury (**Figure 6C – D**). In a parallel treatment group, mice were administered nitrite supplemented drinking water (1.5 g/L) or nitrite-free drinking water for 3 weeks. Nitrite administration increased plasma nitrite levels from 0.79 μM to 6.1 μM in WT mice and from 0.61 μM to 9.5 μM in the SMC-Mfn1-/-mice. Notably, while nitrite significantly decreased intimal hyperplasia in WT mice, it did not have a significant effect in SMC-Mfn1^-/-^ mice (**Figure 6A – B**). Nitrite also decreased Ki67 and YFP double positive cells in WT mice but not in SMC-Mfn1^-/-^ mice (**Figure 6C – D**). Collectively, these data demonstrate that Mfn1 expression prevents SMC proliferation and neointimal hyperplasia in vivo and the nitrite-dependent inhibition of SMC hyperproliferation after injury is dependent on Mfn1 expression *in vivo*.

**Figure 6.**
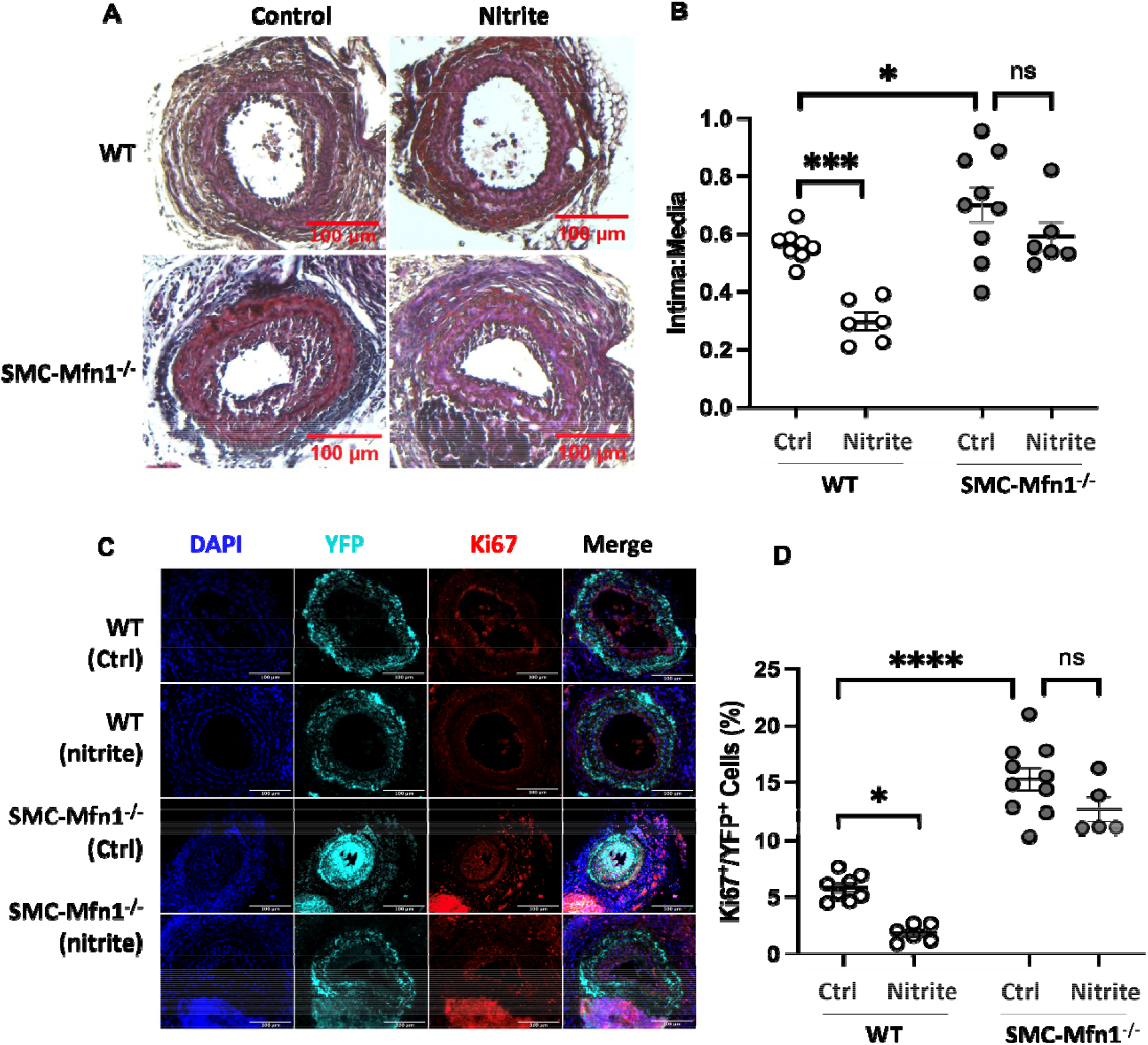
Nitrite-mediated attenuation of vascular remodeling after injury is dependent on Mfn1. **(A)** Images of Masson Trichrome-stained carotid arteries from WT mice and SMC-Mfn1^-/-^ mice subjected to carotid ligation injury followed by administration of normal or nitrite supplemented drinking water for 3 weeks. **(B)** The ratio of intima area to medial area (n=6). **(C)** Representative images of immunofluorescence-stained carotid arteries from WT mice and SMC-Mfn1^-/-^ mice subjected to carotid ligation injury followed by nitrite supplemented drinking water for 3 weeks. Immunofluorescence staining shows nuclei DAPI (nuclei), YFP (SMC tracer) and Ki67 (proliferation marker). **(D)** Quantification of Ki67; YFP double positive cells (n=5). Mean ± SEM; *P<0.05, **P<0.01, ***P<0.001, ****P<0.0001. Mean ± SEM; *P<0.05, **P<0.01, ***P<0.001, ****P<0.0001

## DISCUSSION

The key finding of this study is that endogenous Mfn1 expression maintains VSMC in their differentiated contractile state while loss of Mfn1 expression permits VSMC cell cycle re-entry and proliferation. Further, nitrite treatment increases Mfn1 protein levels, leading to the attenuation of VSMC proliferation *in vitro* and *in vivo*. At a mechanistic level, our data demonstrate that the loss of Mfn1 in VSMC decreases cellular antioxidant capacity which exacerbates cellular ROS levels and in turn propagates cell proliferation. Further, we show that nitrite inhibits degradation of Mfn1 by preventing its ubiquitination.

Our data demonstrate that nitrite selectively upregulates Mfn1 and that Mfn1 regulates VSMC proliferation independent of Mfn2. Mitofusins 1 and 2 have both previously been linked to cell cycle regulation^24-26, 50^. Mitofusin-2 was initially identified as a hyperplasia suppressor gene and was shown to inhibit the p21/ERK/MAP signaling cascade to mediate cell cycle arrest.^50^ More recently, Mfn2 was demonstrated to tether mitochondria to the sarcoplasmic reticulum, facilitating cytosolic calcium removal and supporting mitochondrial energetics for cell cycle stabilization.^26^ It is established that proliferative VSMC show decreased Mfn2 levels compared to quiescent VSMC^51^, that the loss of Mfn2 potentiates pulmonary vascular proliferation^52^, and that overexpression of Mfn2 in balloon angioplasty-injured rat carotid arteries prevents neointima formation.^50^ In contrast to Mfn2, the role of Mfn1 in cell cycle regulation and proliferation is far less studied. Interestingly, emerging studies suggest specific roles for Mfn1 distinct from Mfn2.^29, 30^ For example, a recent study reported Mfn1 but not Mfn2 as a regulator of histone H3 levels in embryonic cells.^53^ This finding is interesting in the context of the prominence of histone activity in controlling VSMC phenotype and proliferation.^43, 54, 55^ Prior studies have also shown that Mfn1 interacts with cyclin B1 during the G2/M phase of the cell cycle, an interaction that propagates mitochondrial fission and cell cycle exit.^48^ However, specific molecular mechanisms by which Mfn1 regulates the cell cycle and proliferation have not been identified. The current study establishes a novel signaling mechanism in which the loss of Mfn1 stimulates oxidant signaling to propagate cell cycle progression and proliferation and identifies a previously unrecognized role of endogenous Mfn1 in maintaining VSMC contractile phenotype.

While the finding that cells deficient in Mfn1 show decreased mitochondrial fusion, depleted ATP and attenuated mitochondrial oxidant production is consistent with prior studies^40, 49^, the loss of antioxidant capacity leading to greater levels of total cellular ROS in Mfn1-deficient cells has previously not been reported. Notably, scavenging of cellular ROS via catalase overexpression significantly attenuates VSMC dysfunction observed with Mfn1 deletion even in the presence of depleted ATP levels. While this appears to be contradictory to published studies demonstrating that mitochondrial dynamics regulate the cell cycle through changes in ATP levels^24^, it is possible that the degree of ATP depletion with Mfn1 deletion is insufficient to disrupt the cell cycle. Further, it is well-established that cellular ROS, particularly from enzymatic sources such as NADPH oxidase, propagate VSMC proliferation through multiple mechanisms including the oxidative activation of key mitogenic kinases or the transactivation of growth factor receptors.^12, 13, 56^ Further studies are required to identify specific targets of ROS in Mfn1-deficient VSMC. Additionally, the mechanism by which Mfn1 maintains catalase and GPx expression is unknown. Our data shows that Mfn1 deletion decreases mtROS production and it is well-established that mitochondrial hydrogen peroxide maintains physiological signaling. Thus, it is interesting to speculate that a decrease in mtROS may mediate signaling to downregulate antioxidant proteins in the cytosolic cellular compartment. Alternatively, a recent study linked Mfn1 deficiency to the modulation of metabolomics as well as alterations in histone acetylation and methylation.^57^ It is possible that Mfn1 metabolically or epigenetically regulates antioxidant gene expression, and this is an area of active ongoing research.

Prior studies have mechanistically connected nitrite-mediated cardiovascular protection to the regulation of mitochondrial dynamics and function^34, 39, 42^. The finding that nitrite-dependent attenuation of intimal hyperplasia is dependent on an increase in Mfn1 levels extends this link. Interestingly, we previously demonstrated that nitrite mediates delayed preconditioning in a cardiomyocyte model of anoxia/reoxygenation via the promotion of mitochondrial fusion due to the inhibition of Drp1 (with no change in Mfn1 levels).^40^ While we did not see any change in Drp1 in the current study, this is potentially due to cell-type specific differences or differential nitrite biochemistry in the delayed preconditioning model. In contrast to prior models in which nitrosative stores may have been partially reduced to nitric oxide in anoxia^40^, the in vitro VSMC data presented in the current study was collected in normoxia and thus presumably independent of hypoxic nitrite reduction. Consistent with a non-reductive mechanism, we observed the nitration of March5. These data are the most recent in a growing body of literature demonstrating nitrite-mediated oxidative post-translational modification of proteins to mediate signaling^41, 58, 59^.

Our in vivo data bridge several pre-clinical studies that demonstrate that nitrite^35-37^ or inhibition of mitochondrial fission^20, 26, 50, 52^ independently attenuate intimal hyperplasia. While genetic modulation of Drp1 and Mfn2 to decrease mitochondrial fusion has shown protective effects in multiple animal models of vascular injury^50, 52^, a pharmacologic strategy to specifically increase mitochondrial fusion that is translatable to humans has not been clearly identified. In contrast, nitrate (which is reduced by an entero-salivary pathway to increase circulating nitrite levels) is well-tolerated in humans and has been shown to decrease vascular dysfunction in multiple human studies.^31, 32, 38, 60-64^ Most relevant to the current study, in the recent NITRATE-OCT trial, dietary nitrate supplementation was shown to significantly decrease in-stent and in-segment late lumen loss after percutaneous coronary intervention compared to placebo treatment.^38^ In separate trials, nitrate supplementation showed a decrease in vascular inflammation as well as attenuation of mitochondrial oxidative stress.^61, 64^ Notably, depletion of Mfn1 has been linked to potentiation of inflammatory signaling in neurons and skeletal muscle.^65, 66^ While mitochondrial morphology has not been measured in clinical trials testing the efficacy of nitrite, the measurement of Mfn1 protein levels after nitrate/nitrite supplementation may be warranted in future studies.

In summary, this study elucidates a novel role for Mfn1 as a key regulator of VSMC proliferation and a specific mechanistic target for nitrite-mediated vascular protection. These findings expand the current understanding of mitofusin biology by distinguishing Mfn1 from Mfn2 in vascular homeostasis and reveal a link between nitrite, mitochondrial dynamics, redox balance, and cell cycle control. Future studies will delineate the molecular mechanisms by which Mfn1 regulates antioxidant gene expression, including potential metabolic or epigenetic pathways, and define how Mfn1 signaling integrates with histone modifications to control VSMC dedifferentiation. The potential to specifically target Mfn1-mediated mitochondrial fusion using nitrite could provide critical insights into novel therapeutic strategies to prevent vascular remodeling not only after systemic vascular injury, but also in other pathologies associated with a dysregulation of mitochondrial dynamics such as pulmonary arterial hypertension and atherosclerosis.

