## Supplemental Info for "NITRITE INCREASES MITOFUSIN-1 LEVELS TO INHIBIT VASCULAR SMOOTH MUSCLE CELL PROLIFERATION AND PREVENT INTIMAL HYPERPLASIA"

| **Protein** | **Vendor** | **Catalog #** | **Species** | **Dilution** |
| --- | --- | --- | --- | --- |
| **Primary Antibodies** | | | | |
| β-actin | Abcam | 8226 | Mouse | 1:10000 |
| α-tubulin | Calbiochem | CP06 | Mouse | 1:5000 |
| Catalase | Cell Signaling | 14097S | Rabbit | 1:1000 |
| CDK2 | Proteintech | 10122-1-AP | Rabbit | 1:500 |
| CDK4 | Proteintech | 11026-1-AP | Rabbit | 1:1000 |
| Cu/Zn SOD | Sigma | 07-403 | Rabbit | 1:500 |
| Cyclin A | Santa Cruz | H-432 | Rabbit | 1:1000 |
| Cyclin D1 | Santa Cruz | A-12 | Mouse | 1:1000 |
| Cyclin E | Santa Cruz | M-20 | Rabbit | 1:1000 |
| Drp1 | Cell Signaling | 8570S | Rabbit | 1:500 |
| Phospho-Drp1 (Ser637) | Cell Signaling | 4867 | Rabbit | 1:1000 |
| Phospho-Drp1 (Ser616) | Invitrogen | PA5-64821 | Rabbit | 1:1000 |
| GPX1 | Proteintech | 29329-1-AP | Rabbit | 1:1000 |
| Lipoxygenase 5 | Abcam | 169755 | Rabbit | 1:1000 |
| March5 | Abcam | 185054 | Rabbit | 1:250 |
| Mfn1 | Abcam | 126575  221661 | Mouse  Rabbit | 1:1000  1:1000 |
| Mfn2 | Abcam | 56889 | Mouse | 1:1000 |
| Mn SOD | Sigma | 06-984 | Rabbit | 1:1000 |
| MYH11 | Proteintech | 21404-1-AP | Rabbit | 1:500 |
| Nitrotyrosine | Abcam | 7048 | Mouse | 1:1000 |
| NOX1 | Abcam | 131088 | Rabbit | 1:1000 |
| NOX4 | Proteintech | 14347 | Rabbit | 1:500 |
| Opa1 | Abcam | 42364 | Rabbit | 1:1000 |
| SM22α | Proteintech | 10493-1-AP | Rabbit | 1:5000 |
| SMA | Abcam | 5694 | Rabbit | 1:5000 |
| Ubiquitin | Cytoskeleton | AUB01 | Mouse | 1:500 |
| **Secondary Antibodies** | | | | |
| Donkey anti-Rabbit (680 nm) | LI-COR | 926-68073 | Donkey | 1:5000 |
| Goat anti-Mouse (800 nm) | LI-COR | 926-32210 | Goat | 1:5000 |

**Table 1: Antibodies for Western Blot Analysis**

**Table 2: Primers for Real-Time PCR with cDNA using SYBR Reagents**

| **Gene** | **5’ (Forward) Primer** | **3’ (Reverse) Primer** |
| --- | --- | --- |
| 18S | TTG ATT AAG TCC CTG CCC TTT GT | CGA TCC GAG GGC CTA ACT A |
| MYH11 | CAG TTG GAC ACT ATG TCA GGG AAA | ATG GAG ACA AAT GCT AAT CAG CC |
| SM22α | GCA TAA GAG GGA GTT CAC AGA CA | GCC TTC CCT TTC TAA CTG ATG ATC |
| TET2 | AGG TTT GGA GAG AAG GGT AAA G | GAC CTC CGA TAC ACC CAT TTA G |
| Myocardin | CTC AGG CAT TAT CGG GAC ATA G | CAT AGG ATG GCT TCC GGA ATA |

**Table 2: Antibodies for Immunofluorescent Staining**

| **Protein** | **Vendor** | **Catalog #** | **Species** | **Dilution** |
| --- | --- | --- | --- | --- |
| **Primary Antibodies** | | | | |
| Ki67 | Abcam | 15580 | Rabbit | 1:1000 |
| SMA | Thermo Fisher | 14-9760-80 | Mouse | 1:500 |
| GFP (for YFP) | Santa Cruz | B-2 | Mouse | 1:200 |
| Tom20 | Proteintech | 11802-1-AP | Rabbit | 5 μg/mL |
| **Secondary Antibodies** | | | | |
| Anti-Rabbit AlexFluor-488 | Thermo Fisher | A21206 | Donkey | 1:1000 |
| Anti-Mouse AlexFluor-546 | Thermo Fisher | A10036 | Donkey | 1:1000 |
| *Acta2-FITC | Sigma Aldrich | F3777 | Mouse | 1:250 |
| Phalloidin Alexa Fluor 488 | Thermo Fisher | A12379 | - | 1:500 |
| Donkey anti-Rabbit Cy3 | Jackson ImmunoResearch | 711-165-152 | Donkey | 1:1000 |

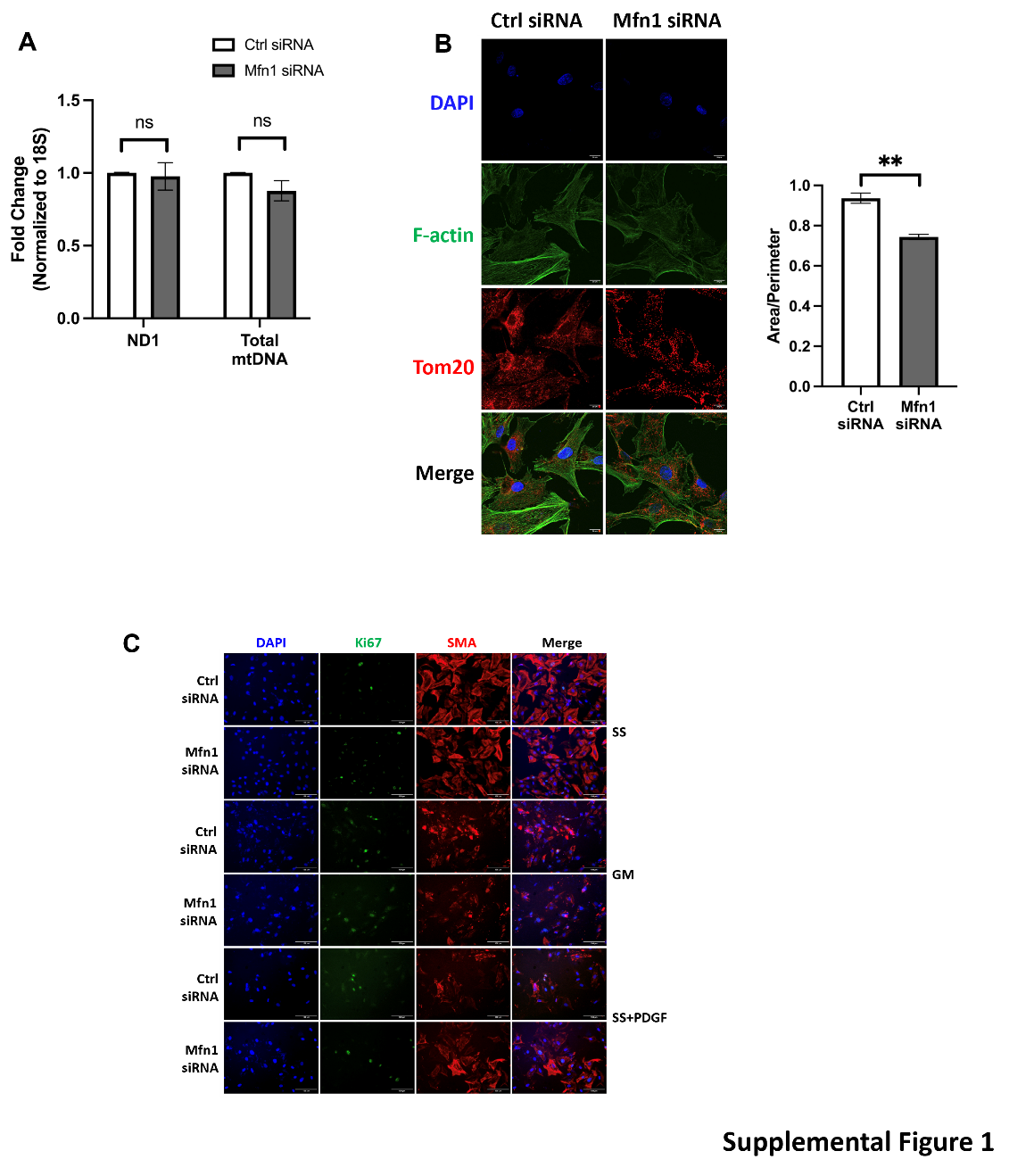

**Supplemental Figure 1: (A)** Relative amount of mRNA for mitochondrial ETC subunit ND1 and total mtDNA levels in RASMC treated with scrambled siRNA (white bars) or Mfn1-targeted siRNA (gray bars). (n=3) **(B)** Representative images of immunofluorescent staining for F-actin (cytoskeletal) and Tom20 (mitochondrial) markers. Images were obtained using Nikon A1 Confocal Microscope System (Nikon) with a 60x oil objective lens and processed and analyzed by NIS-Elements Software (Nikon) and ImageJ and calculation of area/perimeter in RASMC treated with scrambled siRNA or Mfn1-targeted siRNA. **(C)** Representative images of staining for DAPI, Ki67, Smooth Muscle Actin in RASMC grown in growth medium (GM), serum starved (SS) or serum starved then stimulated with PDGF.
